# Spatial turnover of soil viral populations and genotypes overlain by cohesive responses to moisture in grasslands

**DOI:** 10.1101/2022.03.24.485562

**Authors:** Christian Santos-Medellín, Katerina Estera-Molina, Mengting Yuan, Jennifer Pett-Ridge, Mary K. Firestone, Joanne B. Emerson

## Abstract

Although soil viral abundance, diversity, and potential roles in microbial community dynamics and biogeochemical cycling are beginning to be appreciated^1–5^, little is known about the patterns and drivers of soil viral community composition that underlie their contributions to terrestrial ecology. Here, we analyzed 43 soil viromes from a precipitation manipulation experiment in a Mediterranean grassland in California, USA. We recovered 5,315 viral population sequences (vOTUs), and viral community composition exhibited a highly significant distance-decay relationship within the 18 m long field. This pattern was recapitulated in the microheterogeneity of 130 prevalent vOTUs (detected in >=90% of the viromes), which tended to exhibit significant negative correlations between genomic similarity of their predominant allelic variants and distance. Although spatial turnover was also observed in the bacterial and archaeal communities from the same soils, the signal was dampened relative to the viromes, suggesting differences in assembly drivers at local scales for viruses and their microbial hosts and/or differences in the temporal scales captured by viromes and total DNA. Despite the overwhelming spatial signal, vOTUs responsive to a decrease in soil moisture were significantly enriched in a predicted protein-sharing subnetwork of 326 vOTUs linked to 191 known actinobacteriophages, suggesting a genomically cohesive viral response to soil moisture evocative of environmental filtering, potentially by way of actinobacterial hosts. Overall, soil viral ecological processes appear to be highly constrained in space and tightly coupled to the heterogeneous, dynamic soil environment and thus fundamentally different from those of their well-mixed and more thoroughly studied marine counterparts.

## Main text

With an estimated area of 52.5 million square kilometers^6^, grasslands are major contributors to the cycling^7^ and storage^8^ of soil organic carbon at a global scale. Soil microorganisms play key roles in these biogeochemical processes^9, 10^, and, by infecting soil microbiota^11^, viruses likely have substantial direct and indirect impacts on the resulting carbon dynamics^12^. More generally, the potential importance of viruses in soils^1, 2, 13, 14^, together with their measured high abundance (10^7^ to 10^10^ virus-like particles per gram of soil^1^) and improvements in our ability to sequence and track soil viral genomes^12, 15^, has led to a renewed flurry of investigations into soil viral diversity and ecology^3–5, 16–21^. Yet, despite a new appreciation for the vast diversity of soil viruses^3–5, 16–18^, little is known about the factors that govern soil viral community assembly.

To survey dsDNA viral diversity and investigate viral community compositional patterns in Mediterranean grasslands, we collected surface (0 - 15 cm) soil samples from a field site at the Hopland Research and Extension Center in northern California (**Figure 1a**). Since 2017, as part of a large-scale study on the impact of decreased precipitation on soil biotic interactions^22^, experimental plots have received either 100% or 50% of the average historical precipitation at the site via rainfall-excluding shelters and controlled irrigation (**Supplementary Figure 1c,d**). Soils were collected from 22 densely-rooted locations within 15 plots at two time points (March and April, T1 and T2, respectively) during the 2020 growing season of *Avena barbata*, the naturalized annual grass that dominates the site (**Supplementary Figure 1a,b,e**). Soil viral community composition was profiled via 44 viral size-fraction metagenomes (viromes). Assemblies yielded 30,238 contigs >10 Kbp, of which 18,040 were identified as viral by VIBRANT^23^. Viral contig clustering at ≥ 95% average nucleotide identity (ANI) yielded 6,088 approximately species-level^24^ viral operational taxonomic units (vOTUs)^24^ that served as references for read recruitment to establish vOTU relative abundances. After removing vOTUs exclusively detected in single samples and excluding one virome due to poor vOTU recovery, the final dataset consisted of 43 viromes and 5,315 vOTUs.

**Figure 1.**
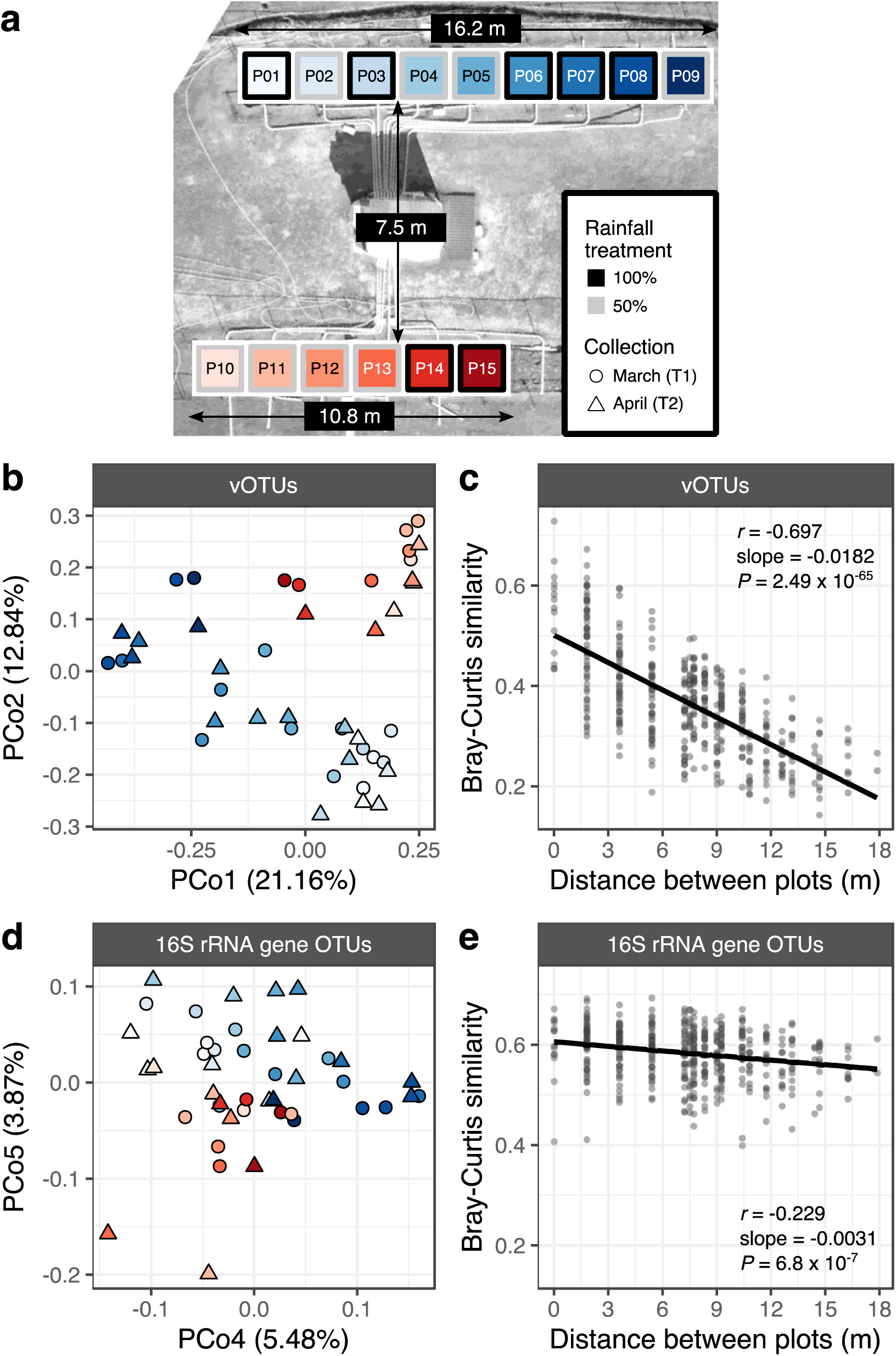
Spatial structuring of viral and prokaryotic communities in a Mediterranean grassland. **(a)** Aerial view of the field site. Colored squares mark the locations of the sampled plots within the upper (blue) and lower (red) blocks. Square outlines indicate the rainfall manipulation regime assigned to each plot. Differences in font color are for legibility only. **(b,d)** Unconstrained analysis of principal coordinates performed on **(b)** vOTU and **(d)** 16S rRNA gene OTU Bray-Curtis dissimilarities. Panel **(b)** displays the first and second axes while panel **(d)** displays the fourth and fifth axes, as they best captured the spatial structuring in **(b)** viral and **(d)** bacterial and archaeal communities. Color reflects the plot from which the sample was collected and corresponds to the gradient palette in panel **(a)**. Point shape represents the collection time point, according to the legend in panel **(a)**. Axis labels indicate the percentage of total variance explained. **(c,e)** Relationship between Bray-Curtis similarity and spatial distance in **(c)** viral communities and **(e)** bacterial and archeal communities. Each point represents a pair of samples, and the spatial distance between them was measured as the length of the line connecting the centers of the corresponding plots. Trend lines display the least squares linear regression model. Inset statistics correspond to the Pearson’s correlation coefficient (*r*), the linear regression slope, and the associated *P*- value.

Viral community beta-diversity patterns were largely explained by the spatial arrangement of plots in the field, as evidenced by a longitudinal gradient captured by the first axis of a principal coordinates analysis (PCoA) and the separation of the upper and lower blocks along the second axis (**Figure 1b**). A significant negative correlation between Bray-Curtis similarity and spatial distance between plots (**Figure 1c**) further revealed that distance-decay relationships were a key driver of viral community composition. These trends were reinforced by substantial differences in individual vOTU detection patterns: of 5,135 vOTUs, 50% were detected in 9 or fewer of the 43 samples (**Supplementary Figure 2b**). Moreover, the percentage of vOTUs shared between pairs of viromes declined steeply as spatial separation increased (**Supplementary Figure 2c**). To assess whether the bacterial and archaeal communities displayed similar spatial patterns, we performed 16S rRNA gene amplicon profiling on total DNA extracted from the same soil samples used to generate the viromes. While some spatial structuring of the bacterial and archaeal communities was observed, this pattern was only evident along the fourth and fifth axes in a principal coordinates analysis (**Figure 2d**). Similarly, even though spatial distance was significantly negatively correlated with microbial community Bray-Curtis similarity (**Figure 2e**), this association was not as pronounced as for the viral communities. In particular, the turnover rate of community similarity over distance (the slope of the distance-decay relationship) for viruses was 5.7 times higher than for bacteria and archaea. These differences in the strength of the spatial patterns between viruses and prokaryotes could be related to differences in the integrated temporal scales captured by DNA pools in viromes compared to total DNA^25^. For example, relic DNA^26^ and DNA from dormant biota^27^ could mask the signal from active microbes in total DNA pools, while the large burst sizes characteristic of viral replication could amplify the signal of recent viral infections in the viromes. The disparity in distance-decay relationships between viral and prokaryotic communities also hints at an oversized role of spatial structuring on the composition of the soil virosphere, an observation that parallels trends recently described for agricultural soils, in which viral but not prokaryotic communities were structured along an 18 m spatial gradient^16^.

**Figure 2.**
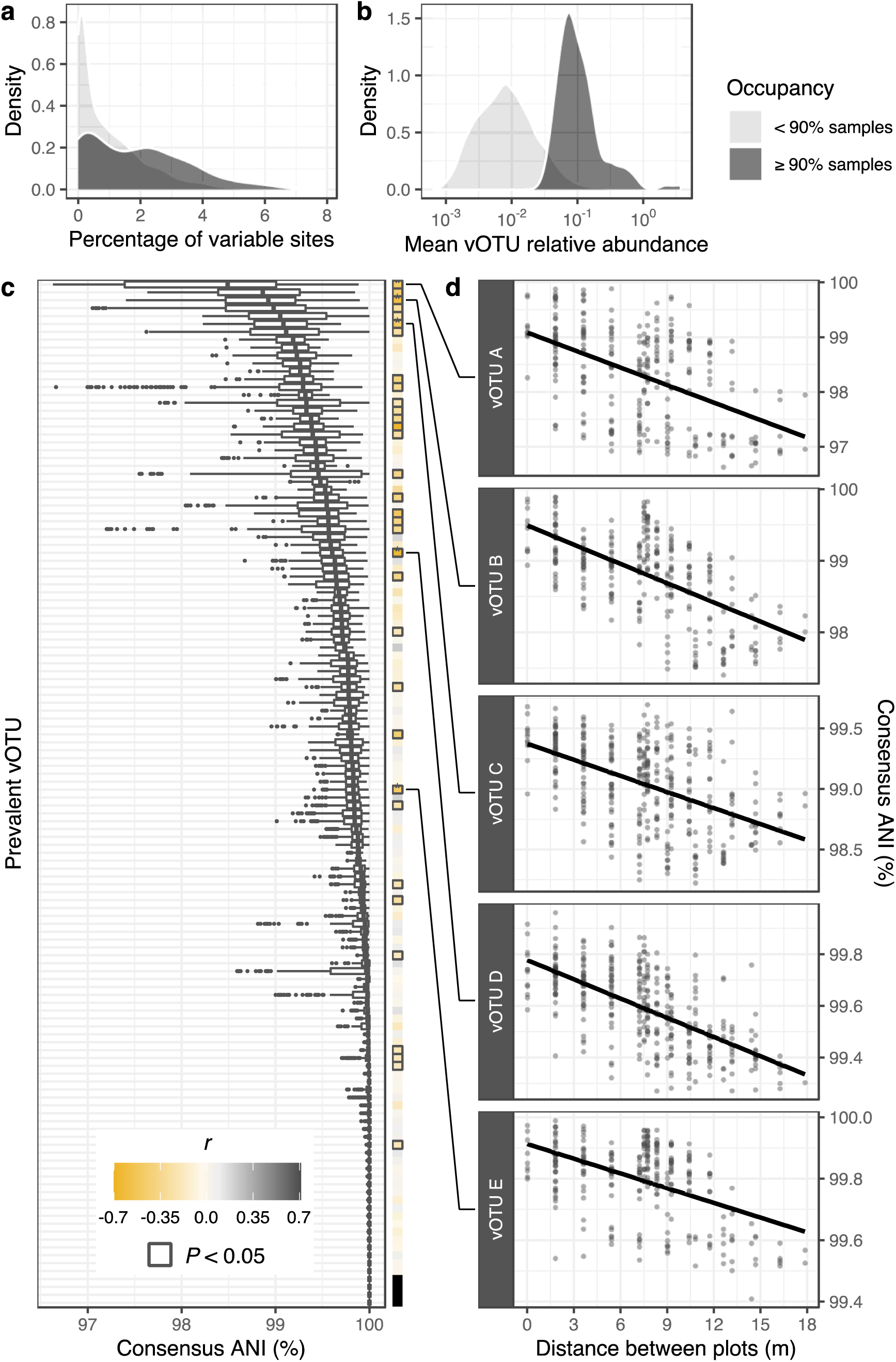
Spatial structuring of viral population microdiversity. **(a,b)** Kernel density plots showing the distributions of **(a)** microdiversity (measured as the percentage of polymorphic sites in a vOTU sequence) and **(b)** mean relative abundance within prevalent (≥ 90% occupancy) and non-prevalent (< 90% occupancy) vOTUs **(c)** Distributions of average nucleotide identities (ANI) for each prevalent vOTU, calculated between pairs of sample-specific vOTU consensus sequences. Each box plot corresponds to a single vOTU, and the y-axis is in rank order (ascending from top to bottom) of the median ANI value for each vOTU. Boxes display the median and interquartile range (IQR), and data points farther than 1.5x IQR from box hinges are plotted as outliers. The heatmap on the right shows the Pearson’s correlation coefficients between consensus ANI and spatial distance. Bold outlines indicate a significant *P-*value (< 0.05) for the correlation after multiple comparisons correction (Holm algorithm). Filled black squares correspond to vOTUs with no variation across samples (i.e., all ANIs were equal to 1). **(d)** The top 5 vOTUs with the most significant correlations (lowest *P*-values) between consensus ANI and spatial distance. Each point represents a pair of samples, and the spatial distance between them was measured as the length of the line connecting the centers of the corresponding plots. The trend line displays the least squares linear regression model. Note that vOTUs are defined in part by sharing >= 95% ANI (see Methods), so within- vOTU ANI values will necessarily be >= 95% ANI. Also note subtle differences in the y- axis range across graphs.

The observed differences in distance-decay patterns for viral and prokaryotic communities suggest that the underlying assembly processes governing spatial structuring at local scales could be differentially impacting these two components of the soil microbiome. For example, contrasting dispersal limitations linked to size, adsorption specificities, and transport mechanisms could result in divergent assembly patterns between viruses and cellular microorganisms^28, 29^. Significant spatial patterns in the abiotic environment (**Supplementary Figure 3a**), including in calcium concentrations (**Supplementary Figure 3b**), are consistent with this possibility as attachment to soil surfaces in the presence of Ca^2+^ can be significantly higher for viruses than for bacteria^29^, potentially impacting their relative movement in soil. At the same time, environmental selection, whereby abiotic and/or biotic factors influence the distribution of microbial populations through selective pressure^30^, could be particularly relevant for viral community assembly, given that viruses depend not only on the edaphic properties that affect their viability and transport but also on the successful infection of suitable hosts^31^. As one example of likely environmental selection on viruses, presumably partly by way of their hosts, RNA viral communities in grasslands were shown to differ significantly in the presence of plant litter and across soil compartments, with patterns similar to those of their host communities^18^. The stronger spatial structuring of viral relative to prokaryotic communities that we observed could be due to a combination of greater dispersal limitation, environmental filtering directly on the viruses, and/or environmental filtering on the hosts, such that even the observed dampened spatial structuring of hosts could amplify the spatial structuring of viruses.

In addition to environmental selection and dispersal, diversification (*i.e.*, the generation of novel genetic variation) can contribute to diversity patterns in microbial communities^30, 32, 33^. To explore the role of spatial structuring on viral genotypic heterogeneity across the field site, we profiled within-population genomic variation. Briefly, using inStrain^34^, we scanned all mapped reads assigned to individual vOTUs and identified polymorphic sites. Then, we reconstructed sample-specific consensus vOTU sequences and assessed inter-sample vOTU genomic similarities via pairwise ANI comparisons. Given that most vOTUs were detected in a limited number of viromes (**Supplementary Figure 2b**), we restricted this analysis to a subset of 130 vOTUs that were detected in at least 90% of the samples. This set of prevalent vOTUs not only tended to display high levels of strain heterogeneity (**Figure 2a**) but also consisted of some of the most abundant viral community members (**Figure 2b**). The ANI distributions revealed a wide range of genomic variation: while some prevalent vOTUs had pairwise similarities close to 0.95 ANI (the threshold used to define a viral population), others appeared nearly clonal across samples (**Figure 2c**). Microdiversity was frequently structured across the longitudinal gradient in the field. For 25% of the prevalent vOTUs and 58% of the 26 most variable such vOTUs (with median ANIs < 99.5%), genomic similarity displayed a significant negative correlation with spatial distance, indicating that the predominant allelic variants tended to diverge with increasing distance (**Figure 2d**). Altogether, these results show that viral community composition and the genetic makeup of viral populations exhibited significant distance-decay relationships across the field, suggesting that most virus-host interactions that lead to successful infections occur within highly localized areas (likely on the scale of meters or less) in these soils.

Although spatial structuring emerged as the predominant driver of viral diversity patterns, we suspected that the contribution of other factors, such as the experimental precipitation treatments, might have been masked by the spatial signal. We thus examined the remaining axes of our principal coordinates analysis (**Figure 1b**) and found that PCo3, which accounted for 8.24% of variance in the dataset, captured a significant distinction between April (time point 2, T2) samples from 50% precipitation plots (‘T2-50’ samples) and the rest of the viromes (**Figure 3a**). Gravimetric soil moisture contents were also significantly lower for these T2-50 samples (**Figure 3b**), reflecting similar water application across all plots shortly before T1 (**Supplementary Figure 1d**) followed by precipitation exclusion treatments predominantly in the 50% treatment plots between the two sampling time points (**Supplementary Figure 1c,d**). The separation of viral communities from the T2-50 samples relative to the rest of the dataset suggests that viral communities were structured more by current or very recent moisture regimes than by historical effects of precipitation treatments that occurred over the preceding three years. We next sought to identify vOTUs with significant responses to these soil moisture patterns so that we could investigate potential commonalities among them. An indicator species analysis revealed 529 vOTUs that were significantly enriched in T2-50 viromes relative to the rest of the samples and 384 vOTUs that were significantly depleted (**Figure 3c**). Much like phylogenetic conservation of some functional traits in bacteria^35^, we wondered whether analyses at higher levels of viral genome conservation might reveal a cohesive response to soil moisture, and we leveraged protein sequence similarity networks to explore this possibility. Using vConTACT2^36^, we constructed a network of vOTUs (nodes), in which each edge indicated significant overlap in predicted protein contents between a pair of vOTUs. We then adapted an algorithm designed to assess local overrepresentation of traits in biological networks^37^ to characterize the network distribution of soil-moisture-responding vOTUs. Briefly, for each vOTU, we identified a local neighborhood of all vOTUs that could be reached, either directly or indirectly, via an edge path with a length shorter than the first percentile of all pairwise node distances in the network. After discarding all local neighborhoods with fewer than 10 vOTUs, we recovered 2,865 subnetworks of highly interconnected nodes with a median size of 39 vOTUs, allowing us to consider many more sizeable groups of related vOTUs than a standard vConTACT2 analysis of ‘genus-level’ viral clusters (VCs)^17, 38^, given that we could only identify 24 VCs with at least 10 vOTUs in this dataset. Next, we performed hypergeometric tests to assess the overrepresentation of vOTUs enriched or depleted in T2-50 samples within each network neighborhood. A total of 108 neighborhoods showed a significant overabundance of vOTUs consistently enriched in T2-50 samples, with 26 to 67**%** of vOTUs in these neighborhoods displaying this trait, compared to only 10**%** of vOTUs in the whole network (**Supplementary Figure 4a-c**). This pattern contrasted with the lack of substantial network aggregation of vOTUs depleted in T2-50 samples, as only four small, local neighborhoods displayed a significant, albeit weak, overrepresentation of this trait (**Supplementary Figure 4a-b**). Interestingly, all of the significantly T2-50 enriched trait neighborhoods were constrained to a single region in the protein-sharing network, indicating that a relatively genomically cohesive group of vOTUs tended to be enriched in T2-50 samples. Furthermore, the indicator vOTUs within this subnetwork covered a range of detection patterns across samples (occupancies) and were distributed across the field (**Supplementary Figure 5a,b**), suggesting that, despite the strong spatial structuring of viral communities overall, this group of genomically related vOTUs responded cohesively to changes in soil moisture, regardless of their locations in the field. To further explore the subnetwork with a significant overrepresentation of T2-50-enriched vOTUs, we performed a second protein-sharing network analysis, this time including all prokaryotic viral genomes in the NCBI RefSeq database. We identified edge connections between vOTUs in the low moisture trait subnetwork and RefSeq viral genomes, in order to assess network neighborhood trends in viral and host taxonomy (**Supplementary Figure 6a-c**). Of 326 vOTUs in the subnetwork, 96 were connected to at least one RefSeq viral genome, all of which were classified as Siphoviridae or as undefined viruses from the order Caudovirales (**Supplementary Figure 6b**), both taxonomic classifications currently under consideration to be replaced by monophyletic genome-based families^39^. More interestingly, all 191 RefSeq viral genomes connected to a trait subnetwork vOTU were isolated from Actinobacteria hosts, suggesting that the low-moisture-responsive vOTU subnetwork was largely comprised of actinobacteriophages (**Figure 3e**). In contrast, only 38% of all 971 vOTUs associated with RefSeq genomes across the entire network were exclusively linked to an actinobacteriophage (**Supplementary Figure 6c)**, indicating a substantial concentration of putative actinobacteriophages in the subnetwork.

**Figure 3.**
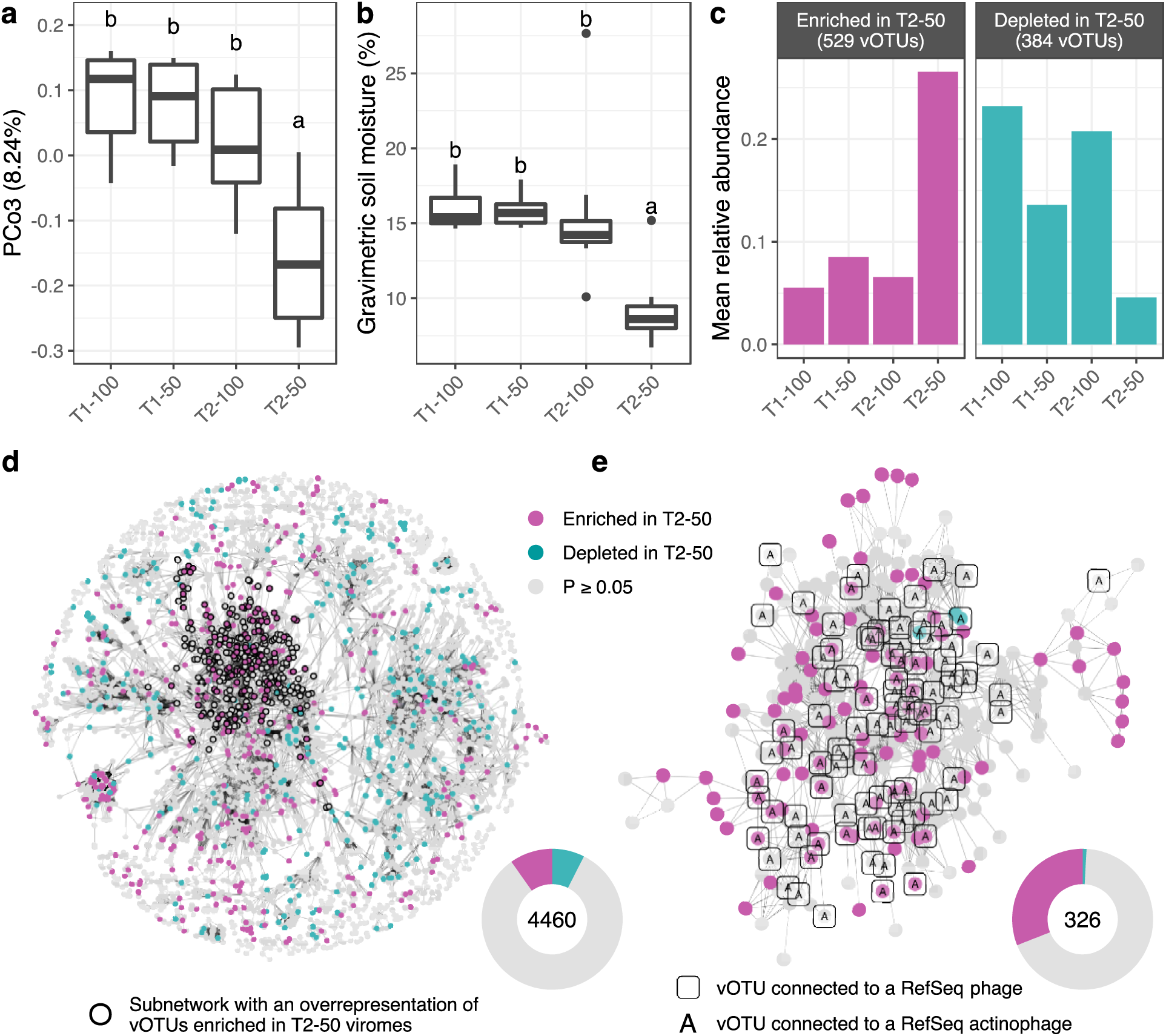
Viral community trends associated with soil moisture content. In **(a)**, **(b)**, and **(c)**, samples are grouped along the x-axis by collection time point (T1 and T2) and precipitation regime (100% and 50%). **(a)** Distribution of scores along the third axis of a principal coordinates analysis performed on vOTU Bray-Curtis dissimilarities. The y-axis label indicates the percentage of total variance explained. The first two axes of the same analysis are shown in Figure 1b. **(b)** Gravimetric soil moisture contents. Boxes display the median and interquartile range (IQR), and data points farther than 1.5x IQR from box hinges are plotted as outliers. In **(a)** and **(b)**, different letters indicate significantly different sample groupings (P < 0.05), as determined by two-tailed Tukey’s range tests. **(c)** Summed mean relative abundances of the sets of vOTUs detected as indicator species differentiating T2-50 communities from the rest of the viromes. Facets distinguish indicator vOTUs that were relatively enriched or depleted, respectively, in T2-50 viromes. **(d)** Gene-sharing network displaying significant overlaps in predicted protein content (edges) between vOTUs (nodes). Node color shows whether a vOTU was an indicator species enriched or depleted in T2-50 samples or not an indicator species (defined by *P*- values below or above 0.05, respectively, from an indicator value permutation test). Bold outlines highlight a subnetwork of all local neighborhoods with a significant overrepresentation of vOTUs enriched in T2-50 viromes (**Supplementary** Figure 4b-c). **(e)** Zoomed in version of the subnetwork highlighted in **(d)**. Nodes surrounded by squares correspond to vOTUs with a significant overlap in their predicted protein contents with any of 971 RefSeq phage genomes, according to the network analysis shown in **Supplementary** Figure 6. All such RefSeq phage genomes with significant links to this subnetwork were from phages isolated on Actinobacteria hosts, indicated by tagging vOTU nodes linked to RefSeq actinophages with the letter “A”. In **(d)** and **(e)**, inset donut plots on the lower right show the total number of vOTUs in the displayed network (center), along with the proportions of the indicator and non-indicator vOTUs in that network (fractions of the circle). Network visualization layouts were generated with the Fruchterman-Reingold algorithm.

Many actinobacteria are drought resistant members of the soil microbiome that can increase their activity and abundance under low moisture conditions across multiple environments^40–43^, including Mediterranean grasslands^44^. While Actinobacteria was among the most abundant phyla in the 16S rRNA gene amplicon profiles, there were no significant differences in its relative abundance across watering treatments or time points (**Supplementary Figure 7a-b**). Furthermore, even though the first axis of a principal coordinates analysis captured a compositional shift from March (T1) to April (T2), there was no clear distinction between T2-50 microbial communities and the rest of the samples. While these results suggest that the effect of low moisture on overall composition observed in the virosphere was not recapitulated by bacterial and archaeal communities (**Supplementary Figure 8**), it is also possible that the presence of genetic material from dead and dormant cells in the total DNA profiles could have concealed underlying ecological dynamics driven by physiologically active microorganisms^26, 27^. In particular, because of its high abundance in soils^45^, extracellular DNA from dead cells can introduce substantial biases in estimates of microbial abundance, especially when the turnover rate of this relic DNA is disrupted by environmental perturbations, such as bacteriophage blooms^25^.

To consider the relic DNA pool in our samples more directly, we recovered reads classified as 16S rRNA gene fragments from virome profiles. Given that viral enrichment in viromes was achieved via 0.22-µm filtration prior to DNA extraction, any bacterial and archaeal sequences present in these libraries likely originated from relic DNA or small (< 0.22 µm) microbial cells^46, 47^. Interestingly, the relative abundance of Actinobacteria 16S rRNA gene reads recovered from T2-50 viromes was significantly higher than in any other group of samples (**Supplementary Figure 7c-d**). This increase in free Actinobacteria DNA, coupled with the enrichment of putative actinophages in the T2-50 subnetworks (**Figure 3e**), suggests a potential increase in the infection and lysis of Actinobacteria hosts under lower moisture conditions.

Overall, our results suggest active and highly dynamic grassland viral communities that are structured over space and can respond cohesively to environmental conditions, such as decreases in soil moisture. The high degree of spatial turnover within one field during one growing season suggests dispersal limitation for most viral populations on scales of meters and months, consistent with estimates that microbes typically travel less than 1 cm per day in most unsaturated soil conditions^48, 49^. Our results suggest that relevant spatiotemporal scales and patterns of viral ecological processes may be fundamentally different in soil from those in well-mixed and more thoroughly studied marine environments. Although evidence for the importance of dispersal in structuring ocean viral communities has been reported^50^, this was on the global scale of ocean currents. In the oceans, many of the same viral populations have been recovered in globally distributed samples and/or in the same place over long periods of time (5 years)^33, 50^. While these scales have yet to be rigorously studied in soil, the stark differences in viral community composition across a single grassland field suggest that we should not expect similar scales of viral population homogeneity. Interestingly, evidence for high turnover in viral genotypes has been shown in the oceans over time^33^, similar to the patterns that we observed here in soil over space. Overall, soil viral community assembly and dispersal patterns seem to be tightly coupled to the heterogeneous and dynamic biotic and abiotic landscape of the local environment, and it will be interesting to see how these local patterns scale over more extensive temporal and spatial distances.

## Materials and methods

### Field experiment and sample collection

Samples were collected as part of a large-scale rainfall manipulation field experiment^22^ at the University of California Hopland Research and Extension Center (39° 00′ 14.6″ N, 123° 05′ 09.1″ W). The field site contained 15 plots (1.8-by-1.8 m) arranged in two separate blocks 7.5 m apart: a 16.2-m-long upper block with 9 plots and a 10.8-m-long lower block with 6 plots (**Figure 1a**). Plot boundaries were delimited by 1-m-deep plastic liners, installed in the spring of 2017, that limited water transfer between adjacent soils. Each plot contained 8 circular subplots (40 cm diameter) delineated by 15-cm-deep PVC collars, and each subplot was further segmented in two halves by a 15-cm-deep plexiglass divider (**Supplementary Figure 1b**). Starting in 2017 and continuing until 2020, plots were exposed to 2 multi-year precipitation regimes with the amount of water received by each plot adjusted to match 100% or 50% of the average historical precipitation at the site. Differential watering was achieved by the periodic deployment of rainfall-excluding shelters (**Supplementary Figure 1c**) and by controlled irrigation of individual plots (**Supplementary Figure 1d**). For this study, soil samples were harvested from 22 subplots distributed across the 15 plots (**Supplementary Figure 1a**). All collected subplots were located within a 60 cm radius from the center of each plot. Collections were performed on March 13th and April 14th, 2020 (T1 and T2, respectively) during the active growth phase of *Avena barbata* (**Supplementary Figure 1e**). For each time point, half of each subplot was destructively harvested. Samples were processed by removing any visible roots, homogenizing the soil, and storing the soil at -80 °C until further processing.

For soil moisture measurements, separate fresh soil sub-samples were collected and processed immediately.

### Virome DNA extraction, library construction, and shotgun sequencing

Due to the COVID-19 2020 lockdown, we could not perform virome extractions on fresh samples as intended and instead stored soils at -80 °C until processing. Soil virions were enriched through filtration and concentration prior to DNA extraction, following a modified version of a previously published protocol^51^. For each sample, 10 grams of soil were resuspended in 10 ml of protein-supplemented phosphate-buffered saline solution (PPBS: 2% bovine serum albumin, 10% phosphate-buffered saline, 1% potassium citrate, and 150 mM MgSO4). To elute virions, soil suspensions were vortexed until homogenized, placed on an orbital shaker (10 min, 400 rpm, 4 °C), and centrifuged (10 min, 3,095 x g, 4°C). Supernatants were recovered and stored briefly at 4 °C, while pellets were resuspended in 10 ml of fresh PPBS for back-extraction of the remaining soil. This process was repeated for a total of three rounds of extraction of the same soil. Supernatants from the same sample were then pooled and centrifuged three times (10 min, 10,000 x g, 4°C), retaining the supernatant and discarding the pellet each time to remove residual soil particles. Purified supernatants were then filtered through a 0.22 µm polyethersulfone membrane to remove cells. Eluted virions in the filtrate were concentrated via ultracentrifugation (2 hrs 25 min, 32,000 x g, 4 °C) in an Optima LE-80K ultracentrifuge with a 50.2 Ti rotor (Beckman-Coulter Life Sciences). Supernatants were removed, and pellets were resuspended in 100 µl of ultrapure water. As previously shown^46, 52^, the DNase treatment step that serves to remove free DNA at this stage is not compatible with samples stored frozen (we suspect that this is because freezing compromises virions), so we were unable to perform a DNase treatment. We have previously shown that non-DNase-treated soil viromes still successfully enrich the viral signal relative to total metagenomes and capture the same ecological trends as DNase-treated viromes from the same samples^46^.

DNA was extracted from the viral fraction with the DNeasy PowerSoil Pro kit (Qiagen), following the manufacturer’s protocol, with the addition of a 10-minute incubation at 65 °C prior to the bead-beating step. Shotgun metagenomic libraries were constructed with the DNA Hyper Prep kit (Kapa Biosystems-Roche), and paired-end sequencing (150 bp) was performed on the NovaSeq S4 platform (Illumina).

### Total DNA extraction, amplicon library construction, and sequencing

Total DNA was extracted from 0.25 g of soil with the DNeasy PowerSoil Pro kit (Qiagen, Hilden, Germany), following the manufacturer’s instructions, with the addition of a 10-minute incubation at 65 °C prior to the bead-beating step. Construction of amplicon libraries followed a previously described dual-indexing strategy^53, 54^. Briefly, universal primers 515F and 806R were used to target the V4 region of the 16S rRNA gene. Amplifications were performed with the Platinum Hot Start PCR Master Mix (ThermoFisher) following the Earth Microbiome Project’s PCR protocol^55^: an initial denaturation step at 94 °C for 3 min, 35 cycles of 94 °C for 45 s, 50 °C for 60 s, and 72 °C for 90 s, and a final extension step at 72 °C for 10 min. Libraries were cleaned with AmpureXP magnetic beads (Beckman Coulter), quantified (Qubit 4 fluorometer), and pooled in equimolar concentrations. Paired-end sequencing (250 bp) was performed on the MiSeq platform (Illumina).

### Soil chemistry and moisture measurements

Soil moisture was calculated as the ratio of mass of water per mass of dry soil. While soil moisture was originally measured for all samples, data for a subset of 11 March samples (5 from 100% plots and 6 from 50% plots) were lost and could not be included in downstream analyses. Soil chemistry profiling was performed by Ward Laboratories (Kearney, NE, USA): soil pH and soluble salts were measured using a 1:1 soil:water suspension; soil organic matter was measured as the percentage weight loss on ignition; nitrate was measured via a KCl extraction; potassium, calcium, magnesium, and sodium were measured via an ammonium acetate extraction; zinc, iron, manganese, and copper were measured via a DTPA extraction; phosphorus was measured via the Olsen method; and sulfate was measured via a Mehlich-3 extraction. Soil chemistry measurements were only performed on the set of 22 soil samples collected in March (T1).

### Bioinformatic processing

#### Virome processing

We used Trimmomatic v0.33^56^ to remove Illumina adapter sequences and quality-trim reads (minimum q-score of 30 evaluated on 4-base sliding windows; minimum read length of 50) and BBDuk v38.82 ^57^ to remove PhiX sequences. Next, we generated *de novo* assemblies of individual libraries with MEGAHIT v1.2.9 ^58^ in meta-large mode (--k-min 27 --k-max 127 --k-step 10), using a contig minimum size threshold of 10,000 bp. Assembled contigs were then classified as viral with VIBRANT v1.2.1^23^ in virome mode. The resulting viral contigs were de-replicated into non-redundant viral operational taxonomic units (vOTUs) with dRep v3.2.2 ^59^, using the following parameters: a threshold of ≥95% average nucleotide identity (ANI) across ≥85% alignment fraction (-sa=0.95, -nc=0.85), single-linkage algorithm for hierarchical clustering (--clusterAlg=single), and filtered nucmer alignments for secondary clustering comparisons (--S_algorithm=ANImf). Representative sequences were selected based exclusively on length (-N50W=0, sizeW=1). Competitive read recruitment against the de-replicated database of vOTUs was performed with bowtie2 v2.4.2 ^60^ in sensitive mode, and the resulting alignments were sorted and indexed with SAMtools v1.11 ^61^. We used CoverM v0.5.0 to generate two vOTU coverage tables: one displaying the trimmed mean coverage (- m=trimmed_mean) and the other displaying the absolute number of mapped reads (- m=count). In both cases, all vOTUs with <75% horizontal coverage were discarded (-- min-covered-fraction=0.75). We filtered out 773 vOTUs that were exclusively detected in single samples and removed one virome due to poor vOTU recovery (136 vOTUs compared to a median of 1,562 vOTUs). The final dataset consisted of 43 viromes and 5,315 vOTUs.

#### Microdiversity profiling

Intrapopulation genetic diversity was characterized with inStrain v1.4.0^34^. First, the bowtie2 alignments described above were parsed with the profile module to identify divergent sites within the set of mapped reads assigned to each vOTU. Variants were only called if a site had a minimum coverage of 5 reads. We then used the compare module to calculate average nucleotide identities between sample-specific consensus sequences, which were reconstructed based on the most common allele detected at each variant site. Pairwise comparisons were considered for downstream analyses only if more than 25% of the vOTU sequence length was covered by the profile module in both samples (percent_genome_compared > 0.25).

#### Gene-sharing network construction

We used Prodigal v2.6.3^62^ in metagenome mode to predict protein content for each de-replicated vOTU and used the resulting amino acid file to construct a gene-sharing network with vConTACT2 v0.9.19 ^36^. The protein alignment step was performed with Diamond^63^, and the protein cluster step was calculated with the MCL algorithm^64^. The NCBI RefSeq database of bacterial and archaeal viral genomes (v85) was included as a reference. Layouts used to visualize the resulting network were calculated with the Fruchterman-Reingold algorithm implemented in the GGally package^65^.

#### Detection and classification of 16S rRNA gene fragments on virome libraries

As previously described^16^, we used SortMeRNA v4.2.0^66^ against representative versions of the bacterial and archaeal SILVA databases^67^ to recover reads containing 16S rRNA gene sequences from the set of quality-filtered virome reads. We assigned taxonomy with the RDP classifier^68^ using the RDP database v18^69^ as reference. A count table was generated from the resulting hierarchical file with the hier2phyloseq() function from the RDPutils package ^70^.

#### Processing of 16S rRNA gene amplicon libraries

Assembly of paired-end reads into single sequences was performed with PANDAseq v2.9^71^, followed by chimeric sequence removal with usearch v6.1^72^. OTU clustering was performed at a 97% sequence identity threshold with the QIIME^73^ implementation of UCLUST v1.2.22^72^ following the open reference protocol against the SILVA database^67^. For consistency with 16S rRNA gene analysis performed on viromes, representative sequences were reannotated with the RDP classifier^68^ using the RDP database v18^69^ as reference. After discarding singletons, the final dataset consisted of 53,854 OTUs.

### Data analysis

All statistical analyses were conducted using R v3.6.3^74^. Unless otherwise noted, all viral analyses were performed on the trimmed mean coverage vOTU table. For vOTU and 16S OTU profiles, Bray-Curtis dissimilarities were calculated on log-transformed relative abundances with the vegdist() function from vegan v2.5-7^75^. To calculate the environmental distance, we first computed the z-score for each soil chemistry variable and then used the dist() function to determine the Euclidean distances between pairs of samples. Principal coordinates analyses were performed with the pcoa() function from ape v5.4-1^76^. Pearson’s correlation tests evaluating the association of spatial distance with Bray-Curtis similarity, community overlap, environmental distance, edaphic variables, and vOTU microdiversity were performed using the cor.test() function with the alternative parameter set to “two.tailed”. The associated linear regression slope was calculated with the lm() function. In all cases, spatial distance between pairs of samples was measured as the length of the line connecting the centers of the corresponding plots. To remove any effect of time point on our spatial correlation analyses, we excluded all pairwise comparisons between samples collected at different time points. For correlation analyses involving multiple comparisons (edaphic variables and microdiversity), p-values were corrected with the Holm algorithm. Indicator species analysis was performed with the multipatt() function from indicspecies v1.7.9 ^77^. For this analysis, we divided the dataset into two groups, one with the T2-50 viromes and the other with the rest of the samples, and we identified vOTUs significantly associated with each group. We used the lm() function to fit linear models evaluating the effect of collection time point and watering treatment on beta-diversity (as captured by individual principal coordinates) and gravimetric soil moisture. We then used the glht() function from the multcomp package^78^ to perform Tukey’s range tests. We used the pairwise.wilcox.test() function to perform pairwise Wilcoxon rank sum tests to assess the effect of collection time point and watering treatment on the relative abundances of Actinobacteria 16S rRNA gene profiles from total DNA and virome DNA. To determine the relative enrichment of vOTUs along the horizontal field transect, we performed a differential abundance analysis with DESeq2^79^, using vOTU non-normalized count tables as input. In particular, we used the DESeq() function to implement negative binomial generalized models to test the effect of the position of each plot on the abundance of individual vOTUs and used the effect size to rank each viral population. All plots were generated with ggplot2 ^80^.

#### Local neighborhood enrichment

To assess whether vOTUs detected as indicator species of T2-50 samples tended to share similar genomic attributes, we adapted a previously described algorithm designed to systematically assess the distribution of traits in biological networks^37^. This algorithm consists of two main steps: (1) for each node in the network, determine a local neighborhood comprised of all nodes that can be directly or indirectly reached via an edge path with a length shorter than a defined threshold; and (2) for each local neighborhood, assess the overrepresentation of a particular attribute among its members. In this study, we used the gene-sharing network generated by vConTACT2^36^, in which nodes represent vOTUs, edges indicate a significant overlap in the predicted content between vOTUs, and edge scores denote the statistical significance of the associated overlap (expressed as - log10 *P*-value). To determine the distance threshold for local neighborhoods, we first calculated the length of the weighted shortest path for each possible pair of nodes in the network and then identified the 1st percentile. We performed this step with the distances() function from the igraph package^81^, using the reciprocal of the edge scores assigned by vConTACT2 as edge weights. We explored the distribution of the following node attributes across the network: (1) coherent enrichment or (2) coherent depletion in T2-50 samples. To assess the overrepresentation of each of these traits in each of the local neighborhoods, we performed hypergeometric tests using the phyper() function with the “lower.tail” parameter set to false. Local neighborhoods with less than 10 nodes were not considered for the overrepresentation analyses. Multiple comparisons correction was performed with the Holm algorithm.

### Data availability

Raw sequences have been deposited in the NCBI Sequence Read Archive under BioProject accession PRJNA818793. Scripts and intermediate files are available at https://github.com/cmsantosm/HoplandViromes.

## Acknowledgments

Most of the work was supported by an award from the U.S. Department of Energy, Office of Science, Office of Biological and Environmental Research, Genomic Science Program, #DE-SC0020163 (grant to MKF, JPR, and JBE). Shotgun metagenomic library construction and high-throughput sequencing were performed by the DNA Technologies and Expression Analysis Core at the UC Davis Genome Center, supported by NIH Shared Instrumentation Grant 1S10OD010786-01.

---

**Supplementary Figure 1.**
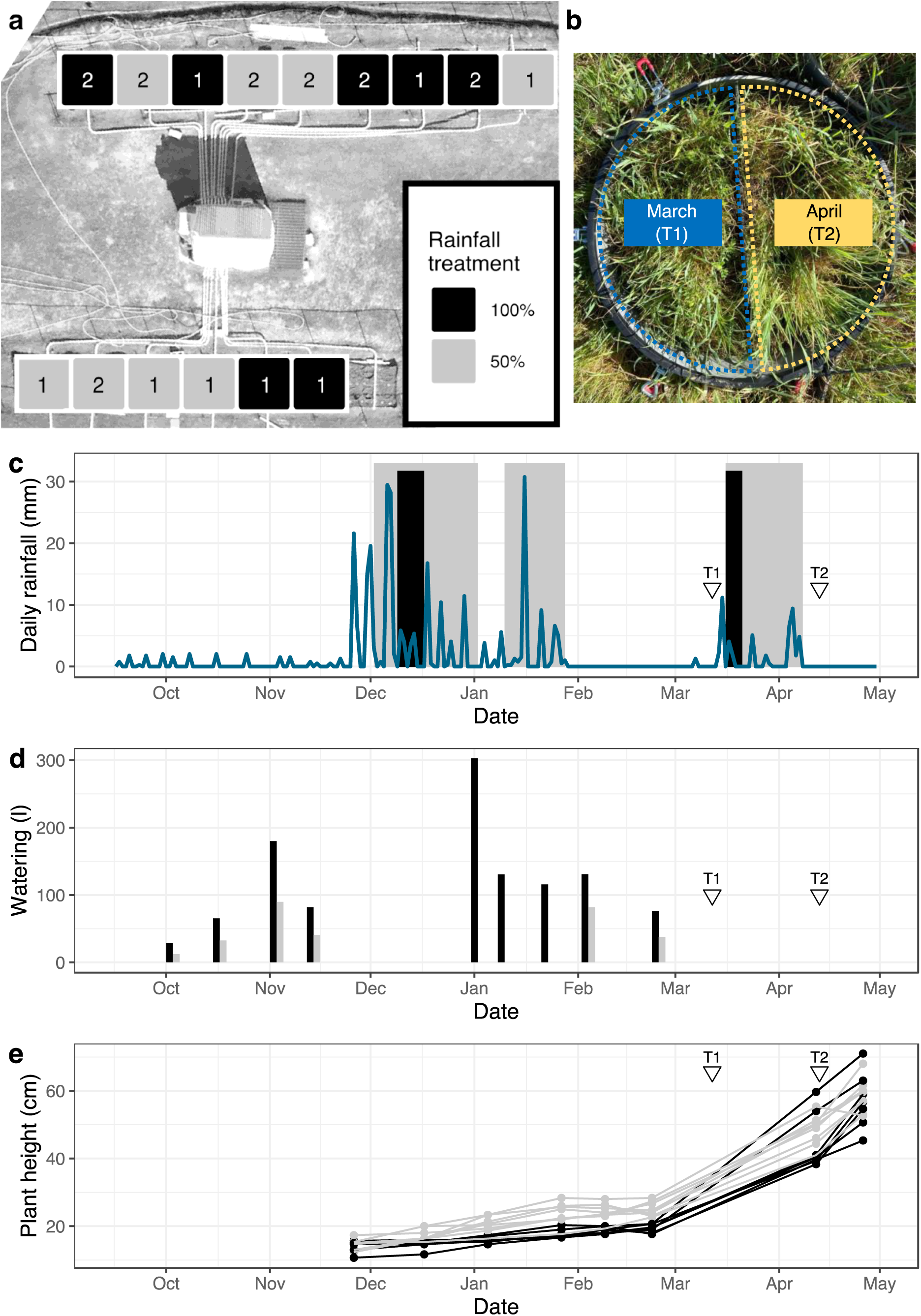
**(a)** Aerial view of the field site. Squares mark the locations of individual plots, and the numbers indicate the number of circular subplots from which samples were collected within each plot. Color indicates the rainfall manipulation treatment for each plot. **(b)** Example of a circular subplot from which samples were collected in this study. Plexiglass dividers segmented subplots into two halves, and one half was destructively harvested at each time point. In panels **(c)** - **(e)**, inverted triangles mark the two collection time points. **(c)** Daily precipitation (blue line) at the field site preceding, during, and shortly after the 2020 sample collection time points (T1 and T2). Background blocks (colored by treatment, as in panel **(a)**) indicate periods when the rainfall-excluding shelters were deployed for each treatment. **(d)** Differential watering events during the months preceding sample collection. Each bar indicates the amount of water added to individual plots through irrigation, based on their assigned watering regime (colored as in panel **(a)**). **(e)** Growth patterns of *Avena barbata* during the 2020 growing season. Each line displays the average height of *A. barbata* in a single plot, colored according to precipitation treatment, as in panel **(a)**.

**Supplementary Figure 2.**
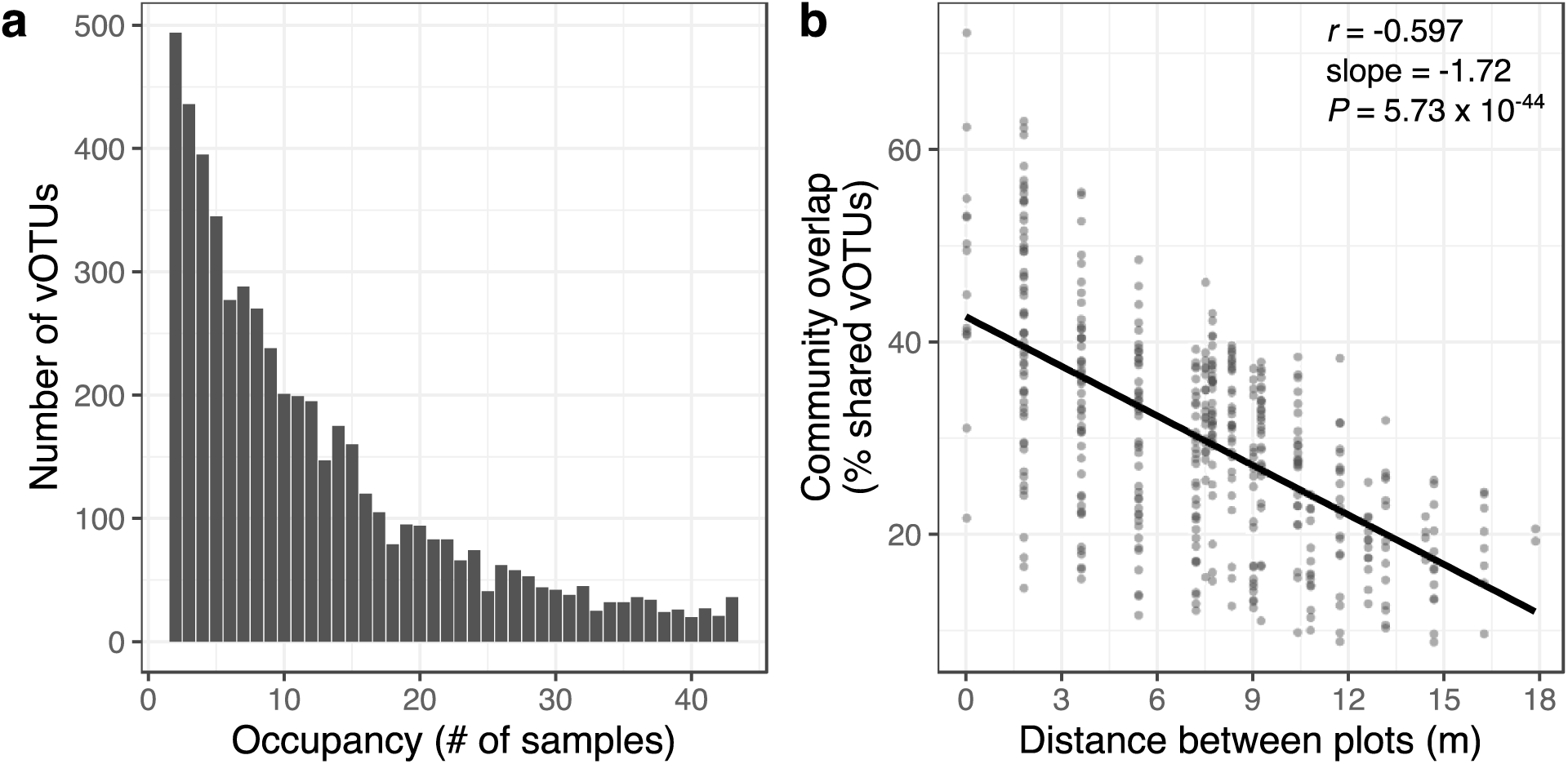
**(a)** Number of vOTUs detected at each occupancy level. **(b)** Relationship between the percentage of vOTUs shared across pairs of samples and spatial distance between plots. Each point represents a pair of samples and the spatial distance between them was measured as the length of the line connecting the centers of the corresponding plots. The trend line displays the least squares linear regression model. Inset statistics correspond to the Pearson’s correlation coefficient (*r*), the linear regression slope, and the associated *P*-value.

**Supplementary Figure 3.**
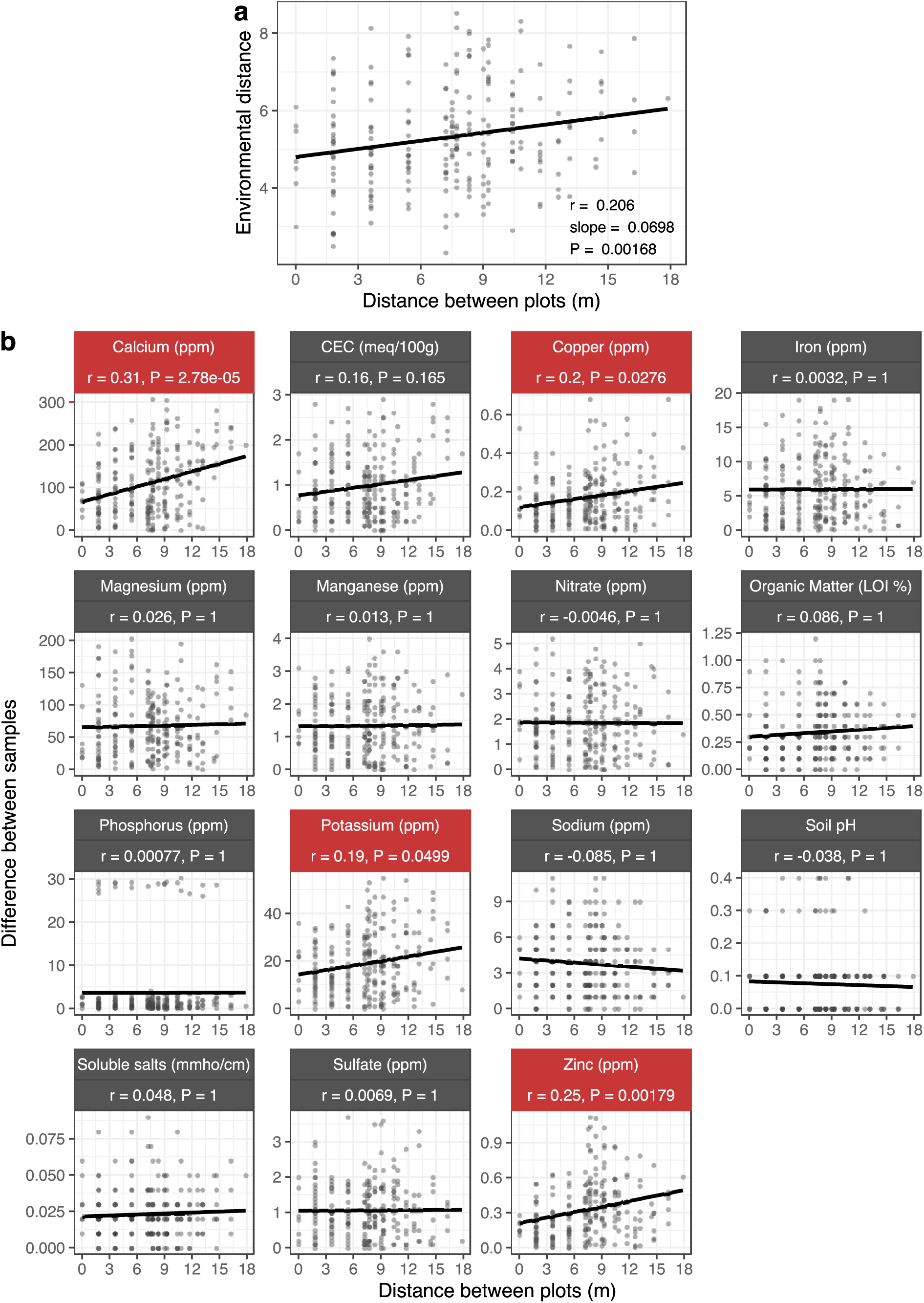
**(a)** Relationship between environmental distance and spatial distance across March (T1) samples, the data subset for which soil abiotic properties were measured. The environmental distance was computed by z-transforming 15 edaphic variables and then calculating their pairwise Euclidean distances. **(b)**. Spatial trends displayed by each of the individual variables used to compute the environmental distance displayed in **(a)**. Each facet corresponds to a single variable with the y-axis indicating the absolute values of the differences between pairs of samples. In both **(a)** and **(b)**, each point represents a pair of samples and the spatial distance between them was measured as the length of the line connecting the centers of the corresponding plots. Trend lines display the least squares linear regression models. Statistics correspond to Pearson’s correlation coefficient (*r*) and associated *P*-value. Variables with a significant correlation in **(b)** are highlighted in red. Abbreviations displayed in the facet names correspond to ppm = parts per million; meq/100g = milliequivalents per 100 grams of soil; LOI% = percent weight loss on ignition; mmho/cm = millimhos per centimeter.

**Supplementary Figure 4.**
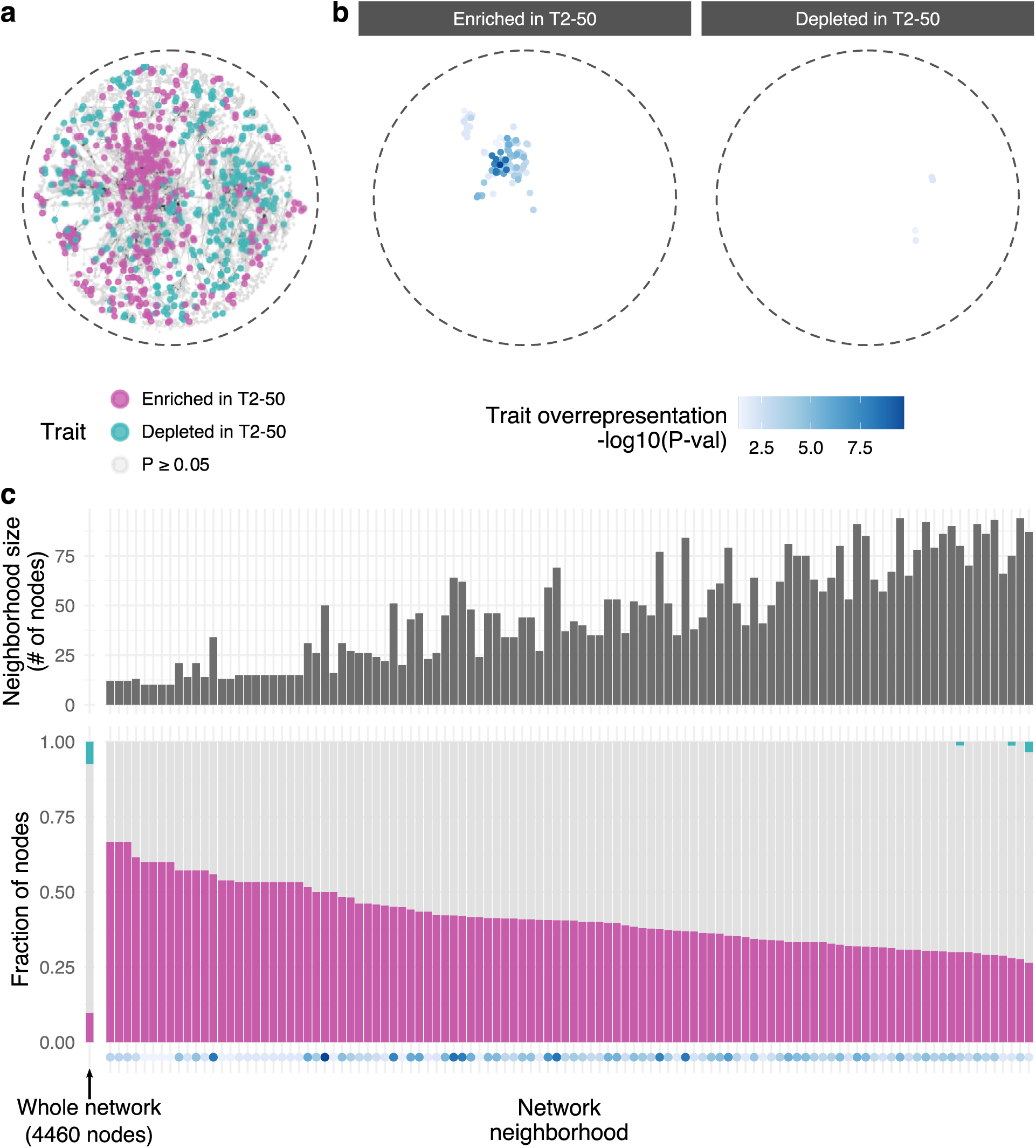
**(a)** Gene-sharing network showing significant overlaps in predicted protein content (edges) between vOTUs (nodes). Node color shows the trait assignment for each vOTU, i.e., whether the vOTU was an indicator species enriched or depleted in T2-50 samples or not an indicator species. **(b)** Distribution of local neighborhoods with a significant overrepresentation of vOTUs enriched (left facet) or depleted (right facet) in T2-50 samples across the network. Each colored point denotes the center of a significant local neighborhood, representing a total of 10 to 94 vOTUs per point. The color gradient indicates the extent of the significance of trait overrepresentation in that neighborhood, with all shades of blue indicating significance and darker shades showing greater significance. Network visualization layout was generated with the Fruchterman-Reingold algorithm. **(c)** Size (upper panel) and trait composition (lower panel) of local neighborhoods with a significant overrepresentation of vOTUs enriched in T2-50 viromes. In the lower panel, each stacked bar plot shows the fraction of indicator vOTUs within a single neighborhood, with the leftmost bar corresponding to the entire network. Dots at the bottom display the statistical significance of the overrepresentation of T2-50-enriched vOTUs in each local neighborhood, following the same color scheme as panel **(b)**.

**Supplementary Figure 5.**
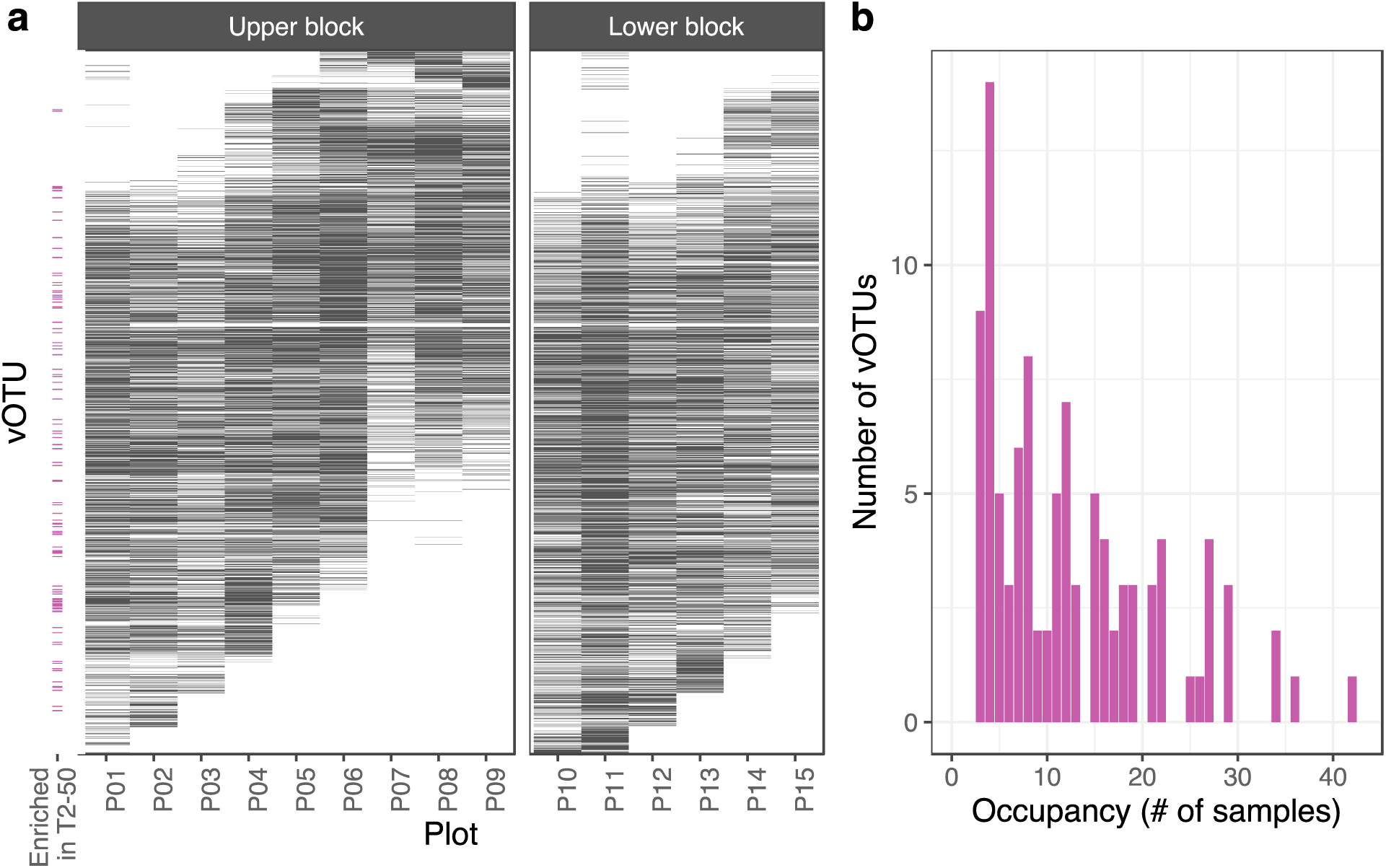
(**a)** Distribution of all 5,315 vOTUs detected in this study across the 15 field plots. Each row represents a single vOTU, and its position along the y-axis is determined by its relative enrichment along the field: vOTUs towards the bottom of the y-axis tended to be more enriched on the North-West (left side of the field in Figure 1a), while vOTUs towards the top tended to be more enriched on the South-East (right side of the field). The pink tick marks on the left side highlight the indicator vOTUs that were enriched in T2-50 samples and that were part of the subnetwork identified in Figure 3e. **(b)** The occupancy spectrum of the indicator vOTUs highlighted in panel **(a)**.

**Supplementary Figure 6.**
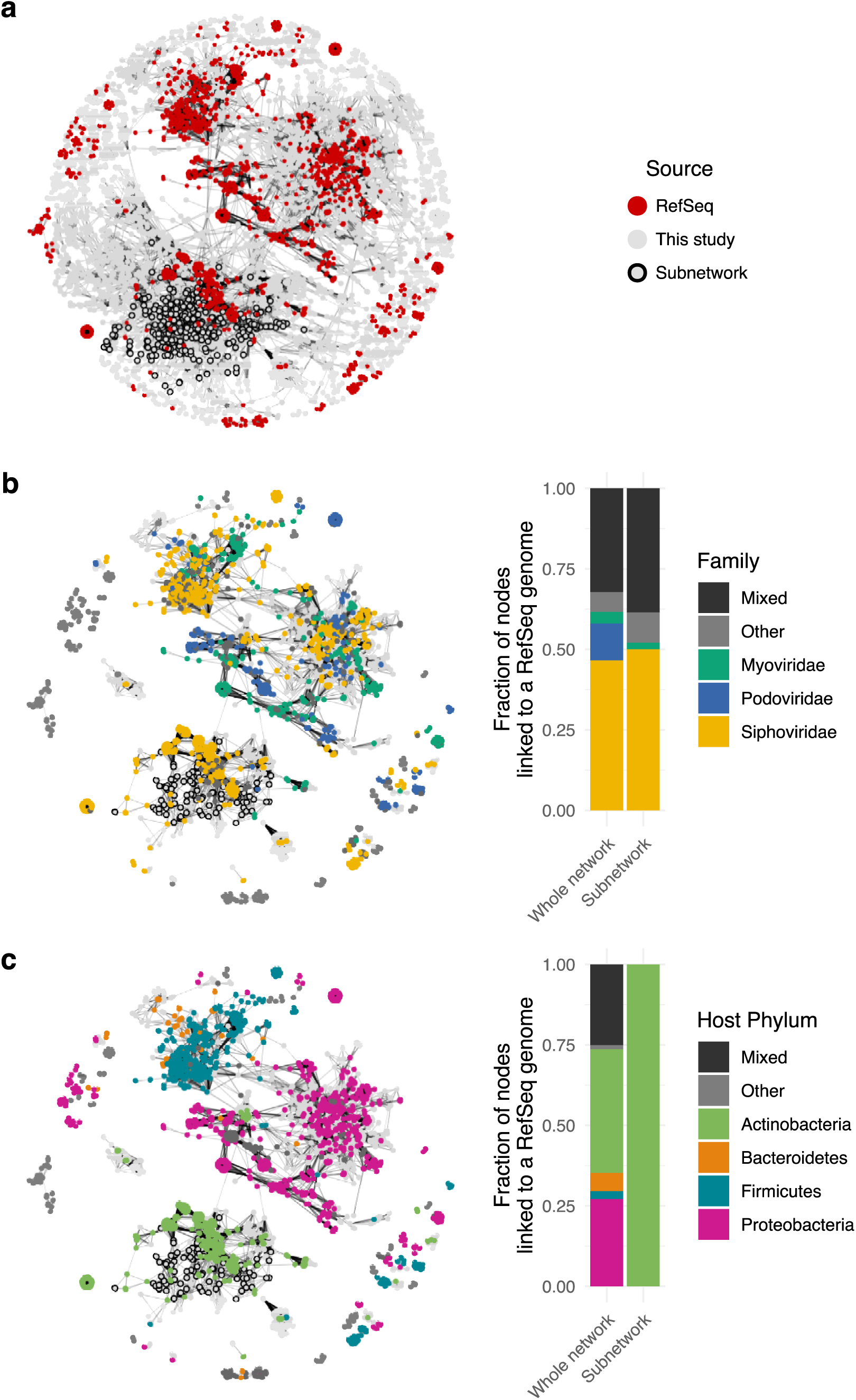
**(a)** Gene-sharing network of vOTUs detected in our study (gray nodes) and prokaryotic virus genomes in RefSeq (red nodes). Edges indicate a significant overlap in the predicted protein content between two viral sequences. The subnetwork highlighted with outlined nodes shows all vOTUs that were part of a local neighborhood with a significant overrepresentation of vOTUs enriched in T2-50 samples (the same nodes outlined in Figure 3d). **(b-c)** The same network, but only showing RefSeq genomes and the subset of vOTUs from our study (light grey) linked by at least one edge to at least one RefSeq genome. Color indicates **(b)** the virus family or **(c)** the host phylum for each RefSeq genome. Accompanying bar plots show the proportion of vOTUs with at least one significant link to a RefSeq genome, with separate bars for the full network and the outlined subnetwork. If a vOTU was linked to multiple RefSeq genomes with differing viral **(b)** or host **(c)** taxonomic classifications, it was labeled as “Mixed”. Network visualization layout was generated with the Fruchterman-Reingold algorithm.

**Supplementary Figure 7.**
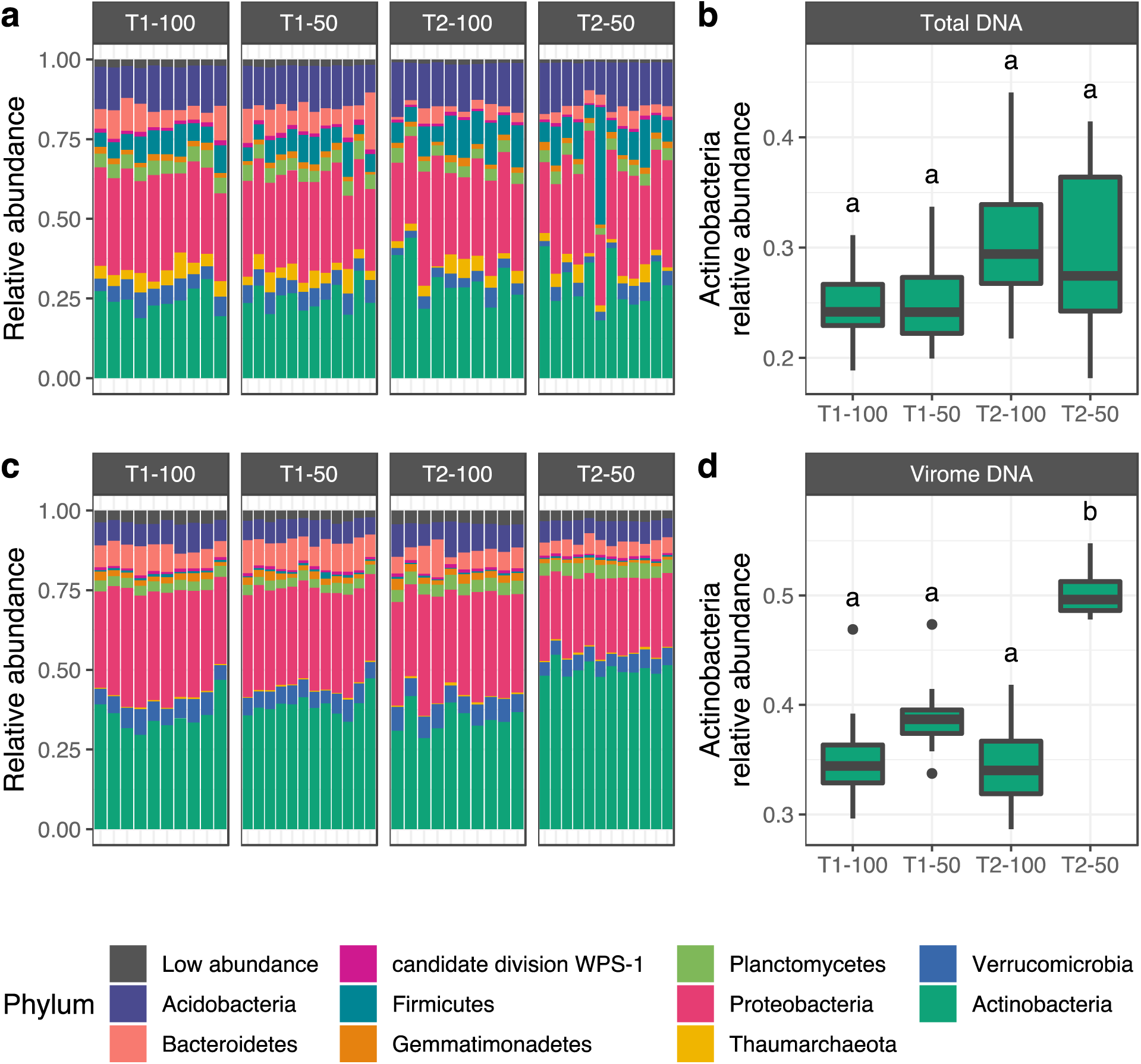
(a,c) Phylum abundances in 16S rRNA gene profiles from **(a)** total DNA 16S rRNA gene amplicon libraries and **(c)** virome DNA libraries. Each stacked bar plot corresponds to a sample, and the 10 most abundant phyla are colored. All other phyla are grouped in the ‘Low abundance’ category. **(b,d)** Relative abundances of Actinobacteria in **(b)** total DNA 16S rRNA gene amplicon libraries and **(d)** virome DNA libraries. Samples are organized by collection time point (T1 and T2) and precipitation treatment regime (100% and 50%). Boxes display the median and interquartile range (IQR), and data points farther than 1.5x IQR from box hinges are plotted as outliers. Letters above boxes indicate significantly different groupings (P < 0.05), as determined by pairwise Wilcoxon’s rank-sum tests.

**Supplementary Figure 8.**
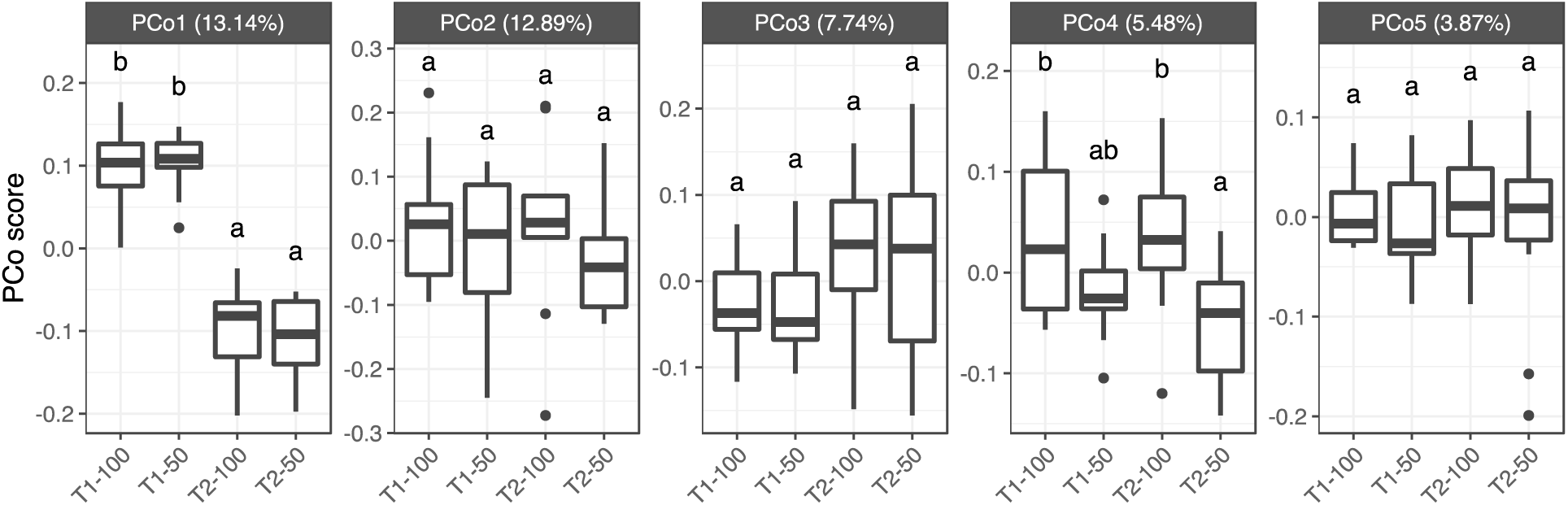
Distribution of scores along the first five axes of a principal coordinates analysis performed on Bray-Curtis dissimilarities from 16S rRNA gene OTU profiles. Samples are organized by collection time point (T1 and T2) and precipitation treatment regime (100% and 50%). Boxes display the median and interquartile range (IQR), and data points farther than 1.5x IQR from box hinges are plotted as outliers. Letters indicate significantly different groupings (P < 0.05), as determined by Tukey’s tests computed for each axis.

## References

1. Williamson, K. E., Fuhrmann, J. J., Wommack, K. E. & Radosevich, M. Viruses in Soil Ecosystems: An Unknown Quantity Within an Unexplored Territory. Annu Rev Virol 4, 201–219 (2017).

2. Kuzyakov, Y. & Mason-Jones, K. Viruses in soil: Nano-scale undead drivers of microbial life, biogeochemical turnover and ecosystem functions. Soil Biol. Biochem. 127, 305–317 (2018).

3. Emerson, J. B. et al. Host-linked soil viral ecology along a permafrost thaw gradient. Nat Microbiol 3, 870–880 (2018).

4. Trubl, G. et al. Active virus-host interactions at sub-freezing temperatures in Arctic peat soil. Microbiome 9, 208 (2021).

5. Schulz, F. et al. Hidden diversity of soil giant viruses. Nat. Commun. 9, 4881 (2018).

6. Food and Agriculture Organization of the United Nations. Grasslands of the World. (Food & Agriculture Org., 2005).

7. Henry, H. A., Gibson, D. J. & Newman, J. A. Biogeochemical cycling in grasslands under climate change. *Grassl*. Clim. Change 1, 115 (2019).

8. Terrer, C. et al. A trade-off between plant and soil carbon storage under elevated CO2. Nature 591, 599–603 (2021).

9. Bardgett, R. D., Freeman, C. & Ostle, N. J. Microbial contributions to climate change through carbon cycle feedbacks. ISME J. 2, 805–814 (2008).

10. Gougoulias, C., Clark, J. M. & Shaw, L. J. The role of soil microbes in the global carbon cycle: tracking the below-ground microbial processing of plant-derived carbon for manipulating carbon dynamics in agricultural systems. J. Sci. Food Agric. 94, 2362–2371 (2014).

11. Starr, E. P., et al. Stable-Isotope-Informed, Genome-Resolved Metagenomics Uncovers Potential Cross-Kingdom Interactions in Rhizosphere Soil. mSphere 6, e0008521 (2021).

12. Emerson, J. B. Soil Viruses: A New Hope. mSystems 4, (2019).

13. Pratama, A. A. & van Elsas, J. D. The ‘Neglected’ Soil Virome - Potential Role and Impact. Trends Microbiol. 26, 649–662 (2018).

14. Fierer, N. Embracing the unknown: disentangling the complexities of the soil microbiome. Nat. Rev. Microbiol. (2017) doi:10.1038/nrmicro.2017.87.

15. Trubl, G., Hyman, P., Roux, S. & Abedon, S. T. Coming-of-Age Characterization of Soil Viruses: A User’s Guide to Virus Isolation, Detection within Metagenomes, and Viromics. Soil Systems 4, 23 (2020).

16. Santos-Medellin, C. et al. Viromes outperform total metagenomes in revealing the spatiotemporal patterns of agricultural soil viral communities. ISME J. (2021) doi:10.1038/s41396-021-00897-y.

17. Ter Horst, A. M., et al. Minnesota peat viromes reveal terrestrial and aquatic niche partitioning for local and global viral populations. Microbiome 9, 233 (2021).

18. Starr, E. P., Nuccio, E. E., Pett-Ridge, J., Banfield, J. F. & Firestone, M. K. Metatranscriptomic reconstruction reveals RNA viruses with the potential to shape carbon cycling in soil. Proc. Natl. Acad. Sci. U. S. A. 116, 25900–25908 (2019).

19. Trubl, G., et al. Soil Viruses Are Underexplored Players in Ecosystem Carbon Processing. mSystems 3, (2018).

20. Van Goethem, M. W., Swenson, T. L., Trubl, G., Roux, S. & Northen, T. R. Characteristics of Wetting-Induced Bacteriophage Blooms in Biological Soil Crust. MBio 10, (2019).

21. Wu, R., et al. DNA Viral Diversity, Abundance, and Functional Potential Vary across Grassland Soils with a Range of Historical Moisture Regimes. MBio 12, e0259521 (2021).

22. Fossum, C. et al. Belowground allocation and dynamics of recently fixed plant carbon in a California annual grassland. Soil Biol. Biochem. 165, 108519 (2022).

23. Kieft, K., Zhou, Z. & Anantharaman, K. VIBRANT: automated recovery, annotation and curation of microbial viruses, and evaluation of viral community function from genomic sequences. Microbiome 8, 90 (2020).

24. Roux, S. et al. Minimum Information about an Uncultivated Virus Genome (MIUViG). Nat. Biotechnol. 37, 29–37 (2019).

25. Lennon, J. T., Muscarella, M. E., Placella, S. A. & Lehmkuhl, B. K. How, When, and Where Relic DNA Affects Microbial Diversity. MBio 9, (2018).

26. Carini, P. et al. Effects of Spatial Variability and Relic DNA Removal on the Detection of Temporal Dynamics in Soil Microbial Communities. MBio 11, (2020).

27. Locey, K. J. et al. Dormancy dampens the microbial distance-decay relationship. Philos. Trans. R. Soc. Lond. B Biol. Sci. 375, 20190243 (2020).

28. Choudoir, M. J. & DeAngelis, K. M. A framework for integrating microbial dispersal modes into soil ecosystem ecology. iScience 103887 (2022).

29. Sasidharan, S. et al. Transport and retention of bacteria and viruses in biochar- amended sand. Sci. Total Environ. 548-549, 100–109 (2016).

30. Nemergut, D. R. et al. Patterns and processes of microbial community assembly. Microbiol. Mol. Biol. Rev. 77, 342–356 (2013).

31. Kimura, M., Jia, Z.-J., Nakayama, N. & Asakawa, S. Ecology of viruses in soils: Past, present and future perspectives. Soil Sci. Plant Nutr. 54, 1–32 (2008).

32. Rubin, B. E. et al. Species- and site-specific genome editing in complex bacterial communities. Nature Microbiology 1–14 (2021).

33. Ignacio-Espinoza, J. C., Ahlgren, N. A. & Fuhrman, J. A. Long-term stability and Red Queen-like strain dynamics in marine viruses. Nat Microbiol (2019) doi:10.1038/s41564-019-0628-x.

34. Olm, M. R. et al. inStrain profiles population microdiversity from metagenomic data and sensitively detects shared microbial strains. Nat. Biotechnol. 39, 727–736 (2021).

35. Martiny, J. B. H., Jones, S. E., Lennon, J. T. & Martiny, A. C. Microbiomes in light of traits: A phylogenetic perspective. Science 350, aac9323 (2015).

36. Bin Jang, H., et al. Taxonomic assignment of uncultivated prokaryotic virus genomes is enabled by gene-sharing networks. Nat. Biotechnol. 37, 632–639 (2019).

37. Baryshnikova, A. Systematic Functional Annotation and Visualization of Biological Networks. Cell Syst 2, 412–421 (2016).

38. Bonilla-Rosso, G., Steiner, T., Wichmann, F., Bexkens, E. & Engel, P. Honey bees harbor a diverse gut virome engaging in nested strain-level interactions with the microbiota. Proc. Natl. Acad. Sci. U. S. A. 117, 7355–7362 (2020).

39. Turner, D., Kropinski, A. M. & Adriaenssens, E. M. A Roadmap for Genome-Based Phage Taxonomy. Viruses 13, (2021).

40. Bouskill, N. J. et al. Pre-exposure to drought increases the resistance of tropical forest soil bacterial communities to extended drought. ISME J. 7, 384–394 (2013).

41. Felsmann, K. et al. Soil bacterial community structure responses to precipitation reduction and forest management in forest ecosystems across Germany. PLoS One 10, e0122539 (2015).

42. Xu, L. et al. Drought delays development of the sorghum root microbiome and enriches for monoderm bacteria. Proc. Natl. Acad. Sci. U. S. A. 115, E4284–E4293 (2018).

43. Santos-Medellín, C. et al. Prolonged drought imparts lasting compositional changes to the rice root microbiome. Nat Plants 7, 1065–1077 (2021).

44. Barnard, R. L., Osborne, C. A. & Firestone, M. K. Responses of soil bacterial and fungal communities to extreme desiccation and rewetting. ISME J. 7, 2229–2241 (2013).

45. Carini, P. et al. Relic DNA is abundant in soil and obscures estimates of soil microbial diversity. Nat Microbiol 2, 16242 (2016).

46. Sorensen, J. W. et al. DNase Treatment Improves Viral Enrichment in Agricultural Soil Viromes. mSystems 6, e0061421 (2021).

47. Nicolas, A. M. et al. Soil Candidate Phyla Radiation Bacteria Encode Components of Aerobic Metabolism and Co-occur with Nanoarchaea in the Rare Biosphere of Rhizosphere Grassland Communities. mSystems 6, e0120520 (2021).

48. Erktan, A., Or, D. & Scheu, S. The physical structure of soil: Determinant and consequence of trophic interactions. Soil Biol. Biochem. 148, 107876 (2020).

49. Tecon, R. & Or, D. Biophysical processes supporting the diversity of microbial life in soil. FEMS Microbiol. Rev. 41, 599–623 (2017).

50. Brum, J. R., et al. Ocean plankton. Patterns and ecological drivers of ocean viral communities. Science 348, 1261498 (2015).

51. Göller, P. C., Haro-Moreno, J. M., Rodriguez-Valera, F., Loessner, M. J. & Gómez-Sanz, E. Uncovering a hidden diversity: optimized protocols for the extraction of dsDNA bacteriophages from soil. Microbiome 8, 17 (2020).

52. Emerson, J. B. et al. Dynamic viral populations in hypersaline systems as revealed by metagenomic assembly. Appl. Environ. Microbiol. 78, 6309–6320 (2012).

53. Caporaso, J. G. et al. Global patterns of 16S rRNA diversity at a depth of millions of sequences per sample. Proc. Natl. Acad. Sci. U. S. A. 108 Suppl 1, 4516–4522 (2011).

54. Edwards, J., Santos-Medellín, C. & Sundaresan, V. Extraction and 16S rRNA Sequence Analysis of Microbiomes Associated with Rice Roots. BIO-PROTOCOL 8, (2018).

55. Thompson, L. R. et al. A communal catalogue reveals Earth’s multiscale microbial diversity. Nature (2017) doi:10.1038/nature24621.

56. Bolger, A. M., Lohse, M. & Usadel, B. Trimmomatic: a flexible trimmer for Illumina sequence data. Bioinformatics 30, 2114–2120 (2014).

57. Bushnell, B. BBTools software package. URL http://sourceforge.net/projects/bbmap (2014).

58. Li, D., Liu, C.-M., Luo, R., Sadakane, K. & Lam, T.-W. MEGAHIT: an ultra-fast single-node solution for large and complex metagenomics assembly via succinct de Bruijn graph. Bioinformatics 31, 1674–1676 (2015).

59. Olm, M. R., Brown, C. T., Brooks, B. & Banfield, J. F. dRep: a tool for fast and accurate genomic comparisons that enables improved genome recovery from metagenomes through de-replication. ISME J. 11, 2864–2868 (2017).

60. Langmead, B., Trapnell, C., Pop, M. & Salzberg, S. L. Ultrafast and memory-efficient alignment of short DNA sequences to the human genome. Genome Biol. 10, R25 (2009).

61. Li, H. et al. The Sequence Alignment/Map format and SAMtools. Bioinformatics 25, 2078–2079 (2009).

62. Hyatt, D. et al. Prodigal: prokaryotic gene recognition and translation initiation site identification. BMC Bioinformatics 11, 119 (2010).

63. Buchfink, B., Xie, C. & Huson, D. H. Fast and sensitive protein alignment using DIAMOND. Nat. Methods 12, 59–60 (2015).

64. van Dongen, S. M. Graph clustering by flow simulation. (2000).

65. Schloerke, B., et al. GGally: Extension to ‘ggplot2’. (2018).

66. Kopylova, E., Noé, L. & Touzet, H. SortMeRNA: fast and accurate filtering of ribosomal RNAs in metatranscriptomic data. Bioinformatics 28, 3211–3217 (2012).

67. Quast, C. et al. The SILVA ribosomal RNA gene database project: improved data processing and web-based tools. Nucleic Acids Res. 41, D590–6 (2013).

68. Wang, Q., Garrity, G. M., Tiedje, J. M. & Cole, J. R. Naive Bayesian classifier for rapid assignment of rRNA sequences into the new bacterial taxonomy. Appl. Environ. Microbiol. 73, 5261–5267 (2007).

69. Cole, J. R. et al. Ribosomal Database Project: data and tools for high throughput rRNA analysis. Nucleic Acids Res. 42, D633–42 (2014).

70. Quensen, J. RDPutils: R Utilities for Processing RDPTool Output. (2018).

71. Masella, A. P., Bartram, A. K., Truszkowski, J. M., Brown, D. G. & Neufeld, J. D. PANDAseq: paired-end assembler for illumina sequences. BMC Bioinformatics 13, 31 (2012).

72. Edgar, R. C. Search and clustering orders of magnitude faster than BLAST. Bioinformatics 26, 2460–2461 (2010).

73. Caporaso, J. G. et al. QIIME allows analysis of high-throughput community sequencing data. Nat. Methods 7, 335–336 (2010).

74. R Core Team. R: A Language and Environment for Statistical Computing. (2018).

75. Oksanen, J. et al. vegan: Community Ecology Package. (2018).

76. Paradis, E., Claude, J. & Strimmer, K. APE: Analyses of Phylogenetics and Evolution in R language. Bioinformatics 20, 289–290 (2004).

77. De Cáceres, M. & Legendre, P. Associations between species and groups of sites: indices and statistical inference. Ecology 90, 3566–3574 (2009).

78. Hothorn, T. et al. Multcomp: simultaneous inference in general parametric models. R package version 1–3 (2014).

79. Love, M. I., Huber, W. & Anders, S. Moderated estimation of fold change and dispersion for RNA-seq data with DESeq2. Genome Biol. 15, 550 (2014).

80. Wickham, H. ggplot2: Elegant Graphics for Data Analysis. (Springer-Verlag New York, 2016).

81. Csardi, G., Nepusz, T. & Others. The igraph software package for complex network research. InterJournal, complex systems 1695, 1–9 (2006).

